# Arc: a therapeutic hub for Alzheimer’s disease

**DOI:** 10.1101/2025.01.16.632170

**Authors:** Antonius M. VanDongen

## Abstract

Alzheimer’s disease (AD) is poised to reach epidemic levels as the world population ages. There is currently no treatment that halts this debilitating disease. Our recent finding that the memory gene Arc regulates the expression of many genes associated with the pathophysiology of AD sets the stage for a new therapeutic approach that is not structurally based on the amyloid hypothesis that has driven most research to date. Neuronal activity-dependent Arc expression is controlled by a chromatin-modification complex containing two enzymes: Tip60 and PHF8. Here, we show that small molecules targeting these proteins inhibit Arc expression. This finding sets the stage for a novel therapeutic approach to combat Alzheimer’s disease. Targeting Arc opens a new frontier of “multi-target” therapy designed to intervene in several aspects of the disease simultaneously. Because of Arc’s role in controlling the expression of multiple genes and pathways implicated in AD, it could serve as a therapeutic hub.

## INTRODUCTION

Our recent discovery [1] that **Arc** (Activity-Regulated, Cytoskeleton-associated protein) controls the expression of many genes implicated in the pathophysiology of Alzheimer’s disease (AD) provides a unique avenue for a novel therapeutic approach. AD is a devastating neurodegenerative disorder characterized by the progressive loss of both synaptic function [2, 3] and long-term memory [4]. The neuron-specific immediate-early gene Arc [5, 6] is a “master regulator” of synaptic plasticity and is required for memory consolidation [7, 8]. Published studies have revealed an association between Arc and AD. A landmark study published in 2011 showed that Arc protein is required to form amyloid (Aβ) plaques [9], a hallmark of AD. Moreover, Arc protein levels are aberrantly regulated in the hippocampus of AD patients [10] and are locally upregulated around amyloid plaques [11]. In contrast, a polymorphism in the Arc gene confers a decreased likelihood of developing AD [12]. It has been shown that spatial memory impairment is associated with dysfunctional Arc expression in the hippocampus of an AD mouse model [12]. These published results suggest that aberrant expression or dysfunction of Arc contributes to the pathophysiology of AD.

Arc was initially localized to glutamatergic synapses, where it regulates AMPA receptor endocytosis, allowing it to regulate synaptic strength. However, it is also found in the neuronal nucleus [13], where it binds beta-spectrin isoform βSpIVΣ5 and is associated with PML bodies, sites of transcription regulation [14]. Nuclear Arc forms a distinct pattern of small puncta that interact with highly dynamic chromatin structures (https://shorturl.at/AFM9w). The nuclear distribution pattern and its association with chromatin and PML bodies suggested that Arc may play a role in epigenetic control of gene expression. Several additional findings supported this idea. Arc interacts with Tip60, a histone-acetyltransferase, and subunit of a chromatin-remodeling complex. Neuronal activity-induced expression of Arc increases endogenous nuclear Tip60 puncta and recruits Tip60 to PML bodies. Arc expression level correlates with the acetylation status of one of Tip60’s substrates: lysine 12 of histone 4 (H4K12), a memory-associated histone mark that declines with age [15]. A survey of histone modifications identified a close association of Arc with H3K27Ac, a marker for active enhancers, and H3K9Ac-S10P, a marker for active transcription [1]. Both these histone modifications, H3K27Ac and H3K9Ac, have recently been shown to be upregulated in late-onset Alzheimer’s disease (AD). These findings suggest that Arc may control long-term memory through epigenetic regulation of gene expression.

To test this idea, we prevented neuronal activity-dependent Arc expression in cultured hippocampal neurons using a short hairpin RNA [1]. Arc knockdown dramatically altered the gene expression program induced by enhanced network activity following chemical long-term potentiation (chem-LTP). RNA-Seq analysis revealed that many genes typically induced by chem-LTP failed to be upregulated when Arc expression was prevented. Another set of genes, ordinarily unaffected by increased network activity, was upregulated when Arc induction was suppressed. A Gene Ontology analysis showed that activity-dependent Arc expression controls the transcription of genes associated with synaptic function, neuronal plasticity, intrinsic excitability, and signaling pathways. Interestingly, about 100 Arc-dependent genes are related to the pathophysiology of AD. Out of 39 AD susceptibility genes identified, 26 were regulated by Arc [1]. The connection between Arc and AD has now come into sharp focus. How can we capitalize on this finding to develop a novel therapeutic strategy for combatting AD? We had earlier discovered that neuronal activity-dependent Arc transcription is epigenetically controlled by a chromatin-modifying complex containing the enzymes Tip60 and PHF8 [16].

Tip60 is a histone deacetylase (HAT) and a binding partner of nuclear Arc, as described above. Studies in a fruit fly AD model displayed an imbalance between Tip60 and HDAC2, with elevated HDAC2 levels leading to the repression of genes associated with learning and memory. Increasing Tip60 levels restored the balance between Tip60 and HDAC2, reinstating normal gene function and improving cognitive abilities [17, 18]. This result suggests that modulating Tip60 activity can counteract neurodegenerative processes linked to Alzheimer’s disease.

PHF8 is the second component of an activity-dependent dual-function chromatin-modifying complex that regulates Arc expression. Mutations in PHF8 cause X-linked intellectual disability and cleft palate. The PHF8-Tip60 complex is rapidly recruited to the Arc promoter in response to neuronal activity, where it facilitates the formation of the transcriptionally permissive histone mark H3K9acS10P by counteracting the repressive mark H3K9me2 [16]. This process enhances Arc transcription. PHF8 (Plant Homeodomain Finger Protein 8) is a histone demethylase already implicated in AD. Studies have shown that decreased PHF8 levels lead to the accumulation of the repressive histone mark H4K20me1, resulting in the upregulation of the mammalian target of rapamycin (mTOR) signaling pathway and subsequent downregulation of autophagy [19]. This disruption impairs the cell’s ability to clear amyloid-beta (Aβ) peptides, leading to their accumulation—a hallmark of AD. In mouse models, PHF8 depletion has been associated with cognitive and neuromotor deficits, further linking its dysfunction to AD-like symptoms. These findings suggest that PHF8 plays a significant role in maintaining neuronal health and that its dysfunction may contribute to the development and progression of Alzheimer’s disease.

The finding that PHF8 and Tip60 together control the expression of Arc highlights the potential of targeting this epigenetic pathway for developing therapeutics for learning and memory disorders, including AD. Therefore, we set out to develop novel drugs targeting these two enzymes and investigated whether they can modulate neuronal activity-dependent Arc expression.

## RESULTS

The findings above highlight the potential of targeting epigenetic regulators like Tip60 and PHF8 in developing therapeutic strategies for Alzheimer’s disease. By restoring the balance between enzymes that modify chromatin structure, it may be possible to alleviate cognitive deficits associated with neurodegenerative conditions. We, therefore, set out to identify small molecules that target Tip60 and PHF8 by an *in-silico* drug screen. High-resolution X-ray crystallographic structures have been published for both enzymes.

Tip60 (HIV-1 <u>T</u>at interacting <u>p</u>rotein, <u>60</u>kDa) is a histone acetyltransferase that transfers acetyl groups to specific histone lysine residues. Its acetyltransferase domain was co-crystalized with its cofactor acetyl-coenzyme A. **Figure 1** illustrates the structure of Tip60 with its bound ligand. Acetyl-CoA is partially buried and occupies a relatively extensive binding site, which looks highly suitable for docking small molecules.

**Figure 1.**
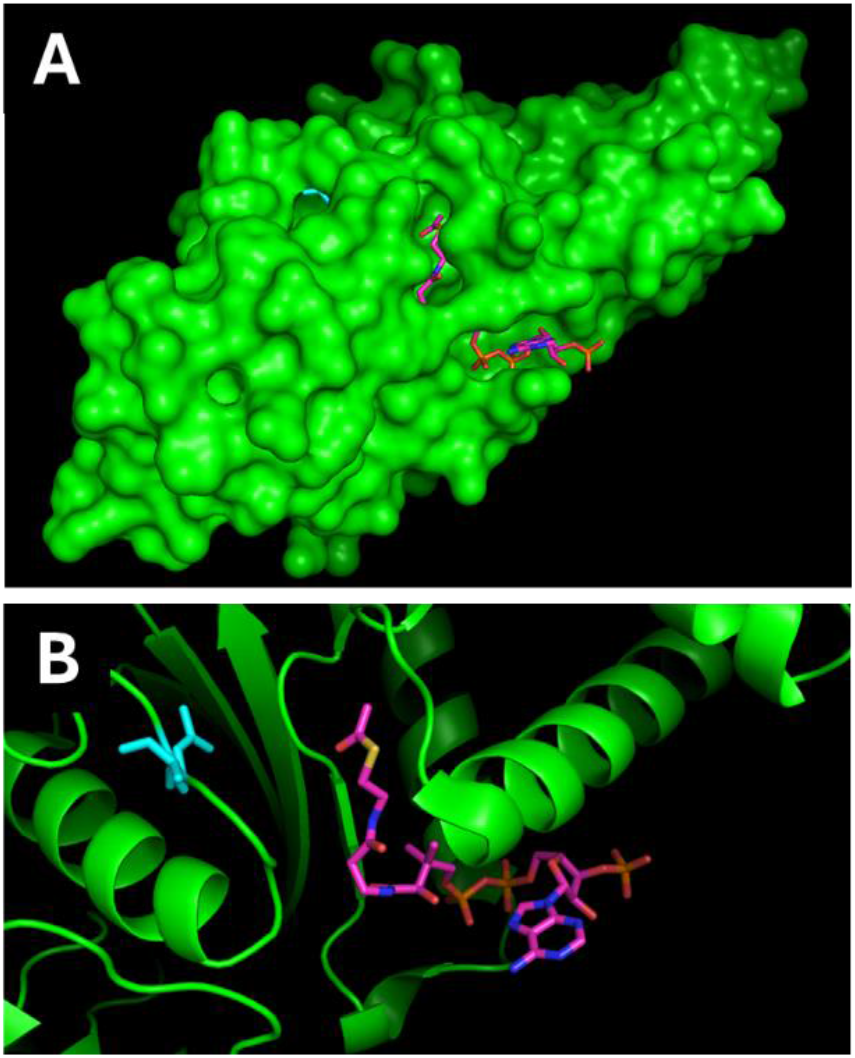
Structure of Tip60. X-ray structure (2ou2.pdb) of the acetyltransferase domain of Tip60, co-crystalized with its cofactor acetyl-CoA. **A**. Tip60 in surface representation (green), acetyl-CoA shown as sticks (magenta). **B**. Zoomed in detail of ligand binding site. Tip60 shown as a cartoon (green), acetyl-CoA as sticks (magenta)

PHF8 (<u>P</u>lant <u>H</u>omeodomain <u>F</u>inger Protein <u>8</u>) is an epigenetic regulator that functions as a histone demethylase. It plays a key role in gene expression and chromatin remodeling. It harbors a Plant Homeo Domain (PHD) that binds Lys4-trimethylated histone 3 (H3K4me3) and a Jumonji C domain that demethylates H3K9me2. See the METHODS section below for additional details. **Figure 2** illustrates the structure of the PHF8-H3-peptide complex. H3K4me3 and H3K9me2 are located in two distinct pockets, which were docked separately.

**Figure 2.**
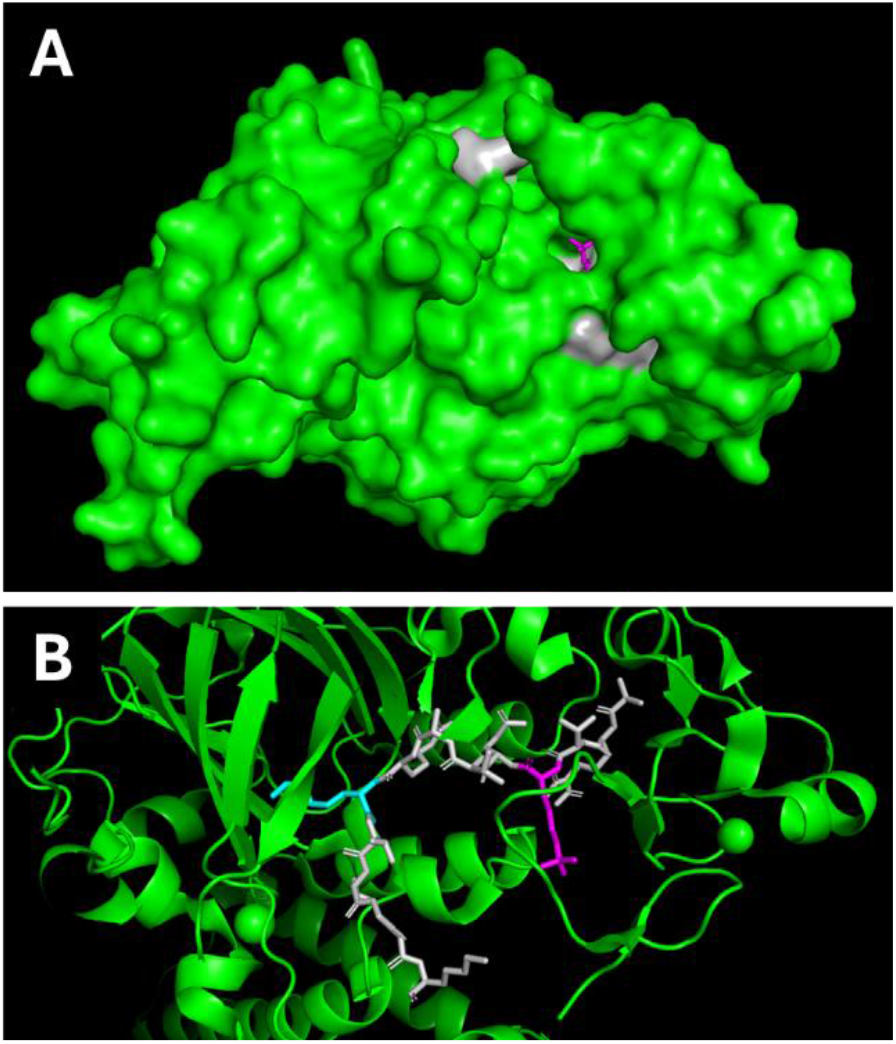
Structure of PHF8. X-ray structure (3KV4.PDB) of PHF8, co-crystalized with a histone H3 peptide. **A**. PHF8 in surface representation (green), H3 peptide in sticks (gray), H3K9me2 in sticks (magenta). **B**. Zoomed in detail of ligand binding site. PHF8 shown as a cartoon (green), H3 peptide as gray sticks, H3K4me3 in magenta, H3K9me2 in cyan.

Small molecules were docked into the three binding sites, Tip60-Acetyl-CoA, PHF8-H3K4me3, and PHF8-H3K9me2, using the **eHiTs** virtual ligand screening software [20-22]. See the METHODS section below for more details. eHiTS is an exhaustive and systematic docking tool with many automated features that simplify the drug design workflow. The optimal 3D pose for each drug in their binding site is determined, and their binding affinity is estimated. A database of 9 million drugs available for purchase from MolPort LLC (www.molport.com) was downloaded from the ZINC repository (www.zinc15.docking.org) [23] in 3D SDF format and screened using eHiTs. Drugs were ranked by predicted affinity, and top-ranking compounds with the highest affinity were ordered for *in vitro* evaluation. **Figure 3** shows the optimal poses of top-ranking drugs in their binding site, next to the original ligand.

**Figure 3.**
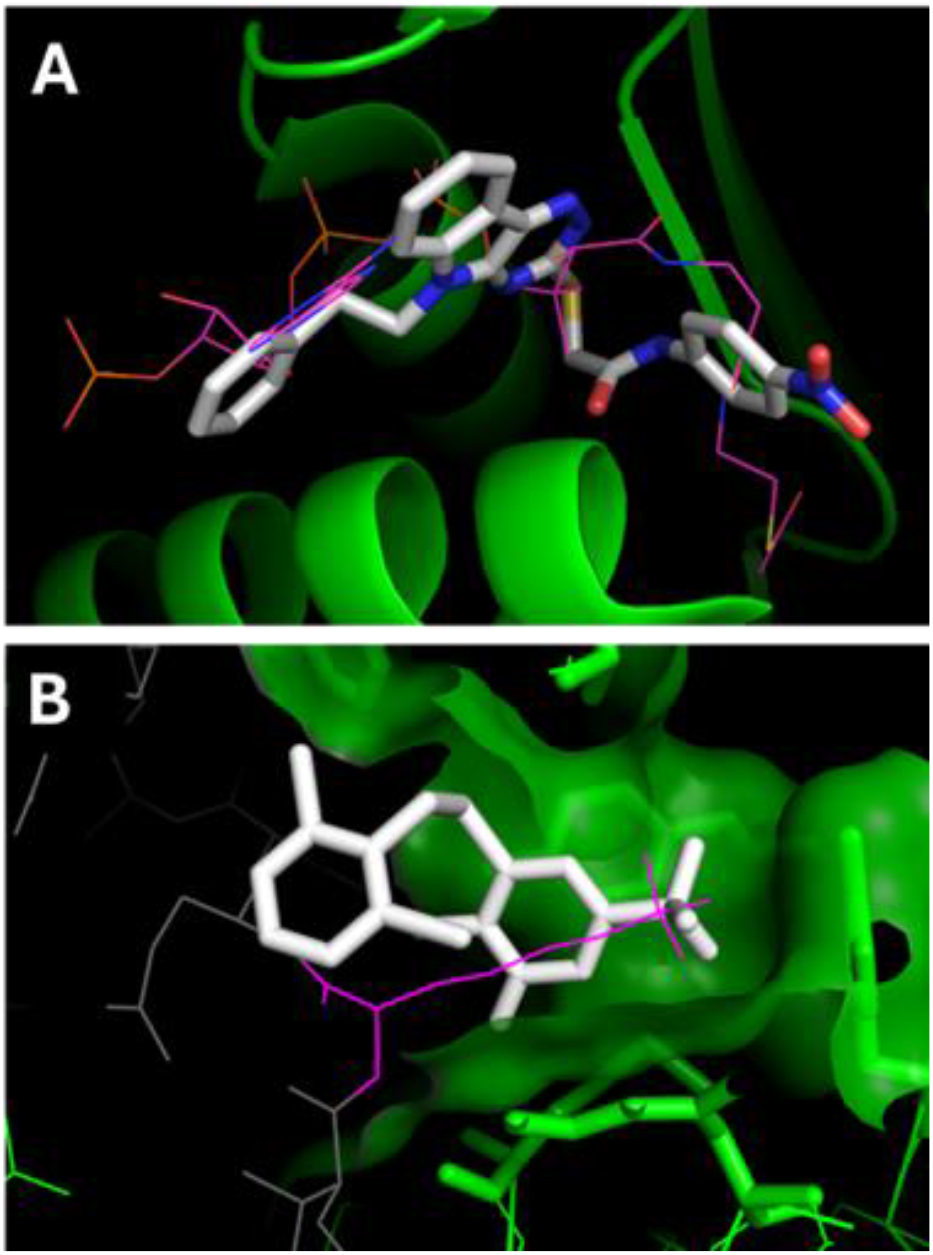
Optimal poses for top-ranking drugs. **A**. Tip60 (green) with drug Tip60.1 (white sticks) and ligand acetyl-CoA (magenta lines). **B**. PHF8 (green) with drug at H3K4me3 (magenta line).

Tip60 and PHF8 are components of a chromatin-modifying complex that controls the neuronal activity-dependent expression of the Arc gene [16]. The small molecules identified in the virtual ligand screen are expected to inhibit the function of these two enzymes by occupying critical binding sites for cofactors and substrates, thereby inhibiting Arc expression. The efficacy of top-ranking drugs was evaluated using cultured hippocampal neurons. In this *in vitro* assay, neurons are plated at a high density (1.5 × 105/cm^2^) in a 96-well culture dish format, forming spontaneously active neuronal networks. At three weeks in culture, neurons have a mature morphology with long axons extending into the network and dendrites covered with spines. The chemical LTP protocol, which combines 4-aminopyridine, bicuculline, and forskolin (4BF), dramatically increases network activity and induces strong Arc protein expression in about half of the neurons [16, 24-26]. Arc expression peaks at four hours after the addition of the 4BF drug cocktail.

Drugs were dissolved in DMSO at a concentration of 1 mM and added 1:1,000 to the culture medium one hour before the start of the Chem LTP protocol. The final drug concentration was 1 µM, and the DMSO concentration was 0.1%. After four hours of Chem LTP, the neurons were fixed and stained for Arc protein using a specific monoclonal antibody (C9, Santa Cruz, Dallas, TX, USA, SC-17839). Arc proteins were then fluorescently labeled using an anti-mouse secondary antibody coupled with Alexa-Fluor 488 (Molecular Probes, Eugene, OR, USA). Fluorescence images were obtained using widefield microscopy, and images obtained were analyzed using NIS Elements AR version 4.1 (Nikon) as previously described [1]. The Arc intensity (green fluorescence) was measured for all neurons in a well, and an average intensity was calculated for each well.

**Figure 4** shows the results of these experiments. Thirty top-ranking drugs targeting Tip60 or PHF8 were evaluated using the in vitro Arc induction assay alongside the negative control DMSO. Fifteen drugs reduced the intensity of Arc by more than 50% at four hours of Chem-LTP, indicating they partially inhibited the activity-induced expression of Arc.

**Figure 4.**
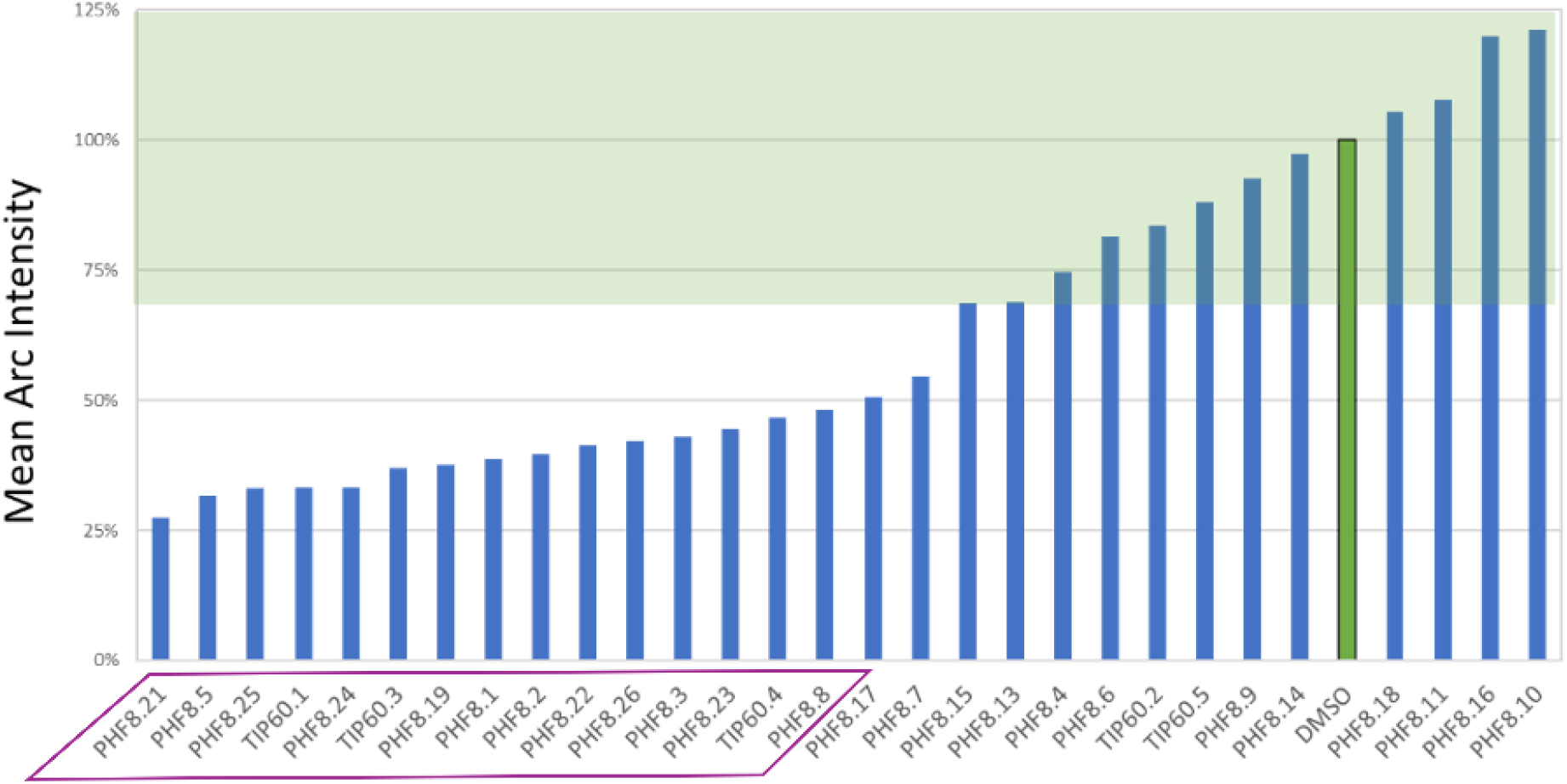
Efficacy of PHF8 and Tip60 drugs. The ability of top-ranking drugs to affect Chem-LTP-induced Arc expression was evaluated using a monoclonal antibody for Arc (C9) and a fluorescent secondary antibody. The bar graph shows the effect of drug treatment on Arc expression. The transparent green bar indicates the range of Arc expression intensities for control DMSO treatments. Fifteen drugs reduced Arc intensity by more then 50%, indicated with a magenta box.

**Figure 5** summarizes the structures of the drugs that showed efficacy in reducing Arc expression by more than 50 %. ZINC IDs are listed for each drug, which can be used to obtain detailed drug information about the drugs at https://zinc15.docking.org/substances. Here, you can also find a list of vendors that sell each drug.

**Figure 5.**
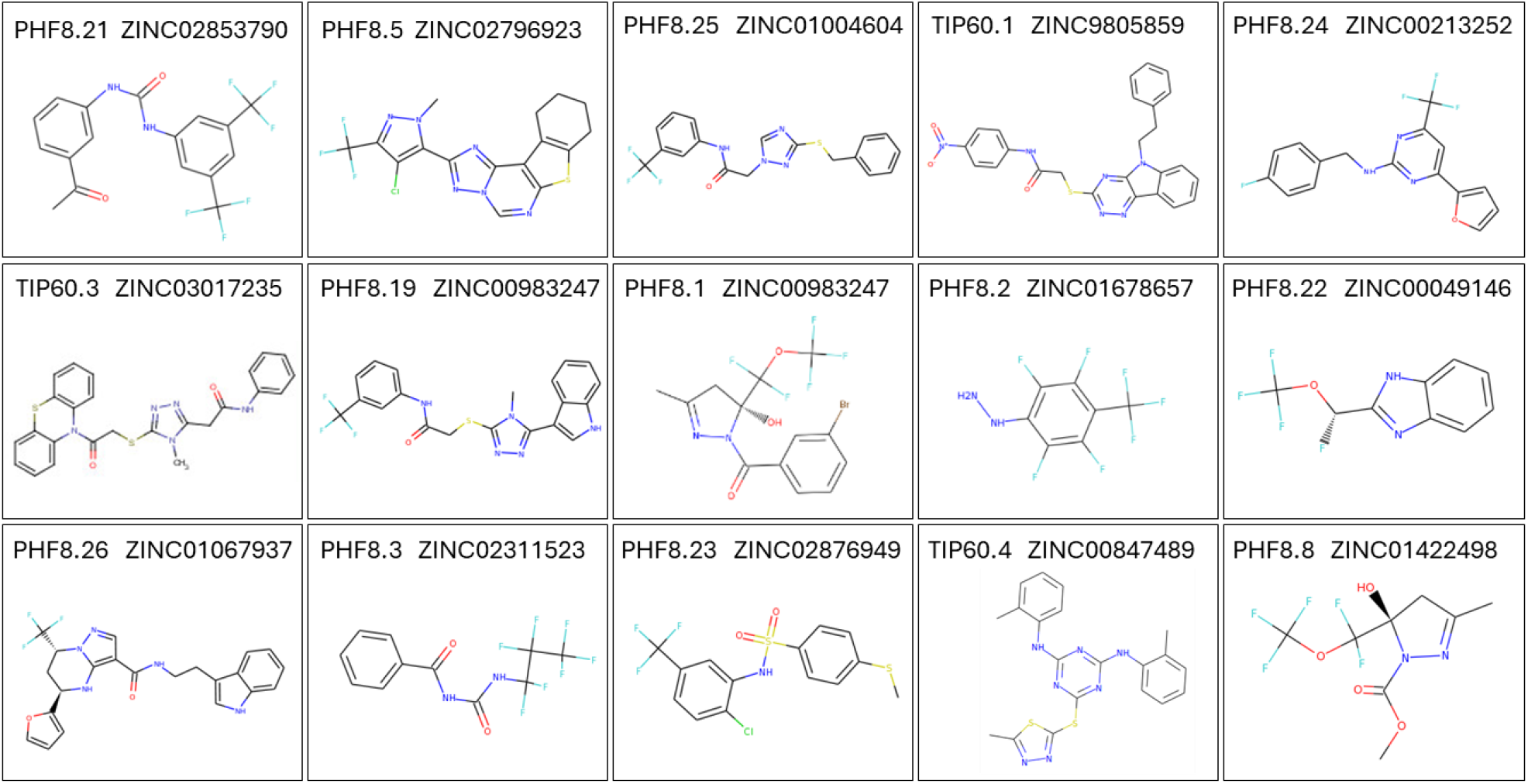
Structures of the drugs that inhibited Arc induction by more than 50%.

## DISCUSSION

In 1906, Alois Alzheimer, a German psychiatrist and neuropathologist, discovered the disease that would later bear his name: Alzheimer’s disease (AD). While studying the brain of a deceased patient named Auguste Deter, who had suffered from severe memory loss, confusion, and unpredictable behavior, Alzheimer observed two distinct pathological features: *amyloid plaques*, clumps of protein fragments, mainly beta-amyloid, that accumulate between nerve cells and 2) *neurofibrillary tangles*, twisted fibers within nerve cells, primarily made up of a protein called tau. These findings marked the first identification of AD as a distinct form of dementia, separate from normal aging. Alzheimer’s discovery provided a foundation for understanding neurodegenerative diseases, though the exact causes and treatment of Alzheimer’s disease are still areas of ongoing research.

Amyloid plaques are a hallmark of Alzheimer’s disease and comprise a protein called beta-amyloid that builds up in the brain. They are a collection of amyloid-beta (Aβ) peptides, strings of amino acids that clump together and form insoluble plaques. The most common form of Aβ peptide in plaques is Aβ42, which has 42 amino acids. Amyloid plaques can disrupt cell function and are associated with other degenerative neuronal elements, microglia, and astrocytes. They are mainly found in the amygdala and hippocampus but can be present throughout the cortex.

Advances in neuroimaging, genetics, and molecular biology have expanded our understanding of the mechanisms underlying Alzheimer’s disease (AD) significantly. But despite heroic efforts to find a cure, there is currently no therapy that prevents, stabilizes, or reverses the progression of this disorder that is poised to take on epidemic proportions as the world ages. Over 500 clinical trials for AD have failed. Ninety-eight unique compounds targeting amyloid plaques were tested in phase II and II trials, and none showed efficacy. Thirty phase III trials recruited thousands of patients and cost hundreds of millions to test drugs developed to prevent the formation or reduce existing amyloid plaques. None showed any efficacy in curtailing AD. Therefore, these neuroanatomical features discovered in 1906 appear to be the proverbial red herring. Although both amyloid plaques and neurofibrillary tangles correlate strongly with the pathophysiology of AD, they may not be <u>causative</u> for the disease. The strict focus on the amyloid hypothesis for developing AD therapies, at the exclusion of other potential mechanisms, is responsible for the current lack of adequate treatment. We need to change course and pursue new avenues.

The data shown here point to a potential alternative to the amyloid hypothesis: the memory gene Arc. Arc regulates transcription of many AD susceptibility genes and over 100 genes implicated in the pathophysiology of AD that cover many signaling pathways, possibly exerting a high level of control over AD development. This finding provides a unique avenue for a novel therapeutic approach, which has come into sharp focus with our finding that two chromatin-modifying enzymes, PHF8 and Tip60, act as a dual-function epigenetic “switch” that regulates Arc gene expression. The function of these two proteins can be manipulated using small molecules, thereby allowing pharmacological control of Arc expression. Targeting Arc through pharmacological intervention may offer a novel, neuroprotective therapeutic strategy for AD, addressing the disease’s complexity by modulating multiple pathological pathways. Targeting Arc opens a new frontier of “multi-target” therapy designed to intervene in several aspects of the disease simultaneously. Because of Arc’s role in controlling the expression of multiple genes and pathways implicated in AD, it could serve as a therapeutic hub.

## METHODS

### *In silico* drug screening

X-ray structure data for Tip60 and PHF8 were downloaded from the Protein Data Bank database (www.rcsb.org) as PDB files. The acetyltransferase domain of Tip60 was co-crystalized with its cofactor acetyl-coenzyme A (www.rcsb.org/structure/2ou2). Drugs were docked into the acetyl-CoA binding site. PHF8 was co-crystalized with a histone 3 (H3) derived peptide containing both lysines 4 and 9: **ART(K4)QTAR(K9)STGGK** [27]. Lysine 4 was trimethylated, and lysine 9 was dimethylated (www.rcsb.org/structure/3kv4). Drugs were docked into the two binding sites for the methylated lysines. Docking was performed using eHiTS, an exhaustive and systematic docking tool with many automated features that simplify the drug design workflow [20-22]. The optimal 3D pose for each drug in their binding site is determined, and their binding affinity is estimated. A database of 9 million drugs available for purchase from MolPort LLC (www.molport.com) was downloaded from the ZINC repository (www.zinc15.docking.org) [23] in 3D SDF format and screened using eHiTs. Drugs were ranked by predicted affinity, and thirty top-ranking compounds with the highest affinity were ordered for *in vitro* evaluation.

### Culturing hippocampal neurons

Hippocampi were dissected from the E18 embryos of Sprague Dawley rats and dissociated using papain following the protocol from the Papain Dissociation System (Worthington Biochemical Corporation). Gentle mechanical trituration was performed to ensure complete dissociation of tissues. Dissociated cells were plated on poly-D-lysine coated 96-well plates at a plating density of 1.5 × 10^5^/cm^2^ in Neurobasal medium (Gibco) supplemented with 10% (v/v) fetal-bovine serum (FBS, Sigma), 1% (v/v) penicillin-streptomycin (P/S, Gibco) and 2% (v/v) B27 supplement (Gibco) for 2 hours. FBS-containing medium was then removed and replaced with FBS-free medium, and cells were cultured FBS-free subsequently to prevent astrocytic over-growth. The medium was changed on Days In Vitro (DIV) 5. Subsequently, the medium was changed every three to four days. Experiments were carried out on DIV 18-22.

### Chemical LTP and immunofluorescence

Hippocampal neuronal cultures were treated with a combination of 100 µM 4-aminopyridine (4AP), 50 µM bicuculline (Bic), and 50 µM forskolin for four hours to induce chemical LTP. This drug combination will be referred to as **4BF** henceforth. At the end of the treatment, cells were fixed with 100 % ice-cold methanol at -20°C for 10 min. Cells were washed three times with 1x Phosphate Buffered Saline (PBS, in mM: 137 NaCl, 2.7 KCl, and 12 phosphate buffer) containing 0.1% (v/v) Triton X-100 (PBS-Tx). Cells were washed three times in 1x PBS-Tx and blocked in 2% (w/v) Bovine Serum Albumin (BSA) in 1x PBS for 1 hour at room temperature. Cells were stained for Arc protein with primary anti-Arc antibody (1:300, Santa Cruz, sc-17839) in antibody dilution buffer (1x PBS containing 1% (w/v) BSA, 5% (v/v) serum and 0.05% (v/v) Triton X-100) for 1 hour at room temperature. Cells were washed three times in 1x PBS-Tx. Cells were then probed with 1:1000 AlexFluor 488 (Molecular Probes) for 1 hour at room temperature. Cells were washed thrice, followed by DNA staining with 50 µM DAPI for 20 min at room temperature. Cells were mounted in FluorSave (Calbiochem).

### Widefield microscopy

Fluorescence images were obtained using widefield microscopy, as detailed in Oey et al. (2015) [16]. Images obtained were analyzed using NIS Elements AR version 4.1 (Nikon) to perform background subtraction. Out-of-focus fluorescence was removed using 3D deconvolution (AutoQuant, Media Cybernetics). The Region-Of-Interest (ROI) analysis tool was used to mark nuclei based on DAPI intensity. Each nucleus’s corresponding mean Arc intensity was measured using the automated measurement module. The average mean Arc intensity for all neurons was obtained for each well in the multi-well plates.

